# Acute high-fat high-sugar diet rapidly increases blood-brain barrier permeability in mice

**DOI:** 10.1101/2024.04.14.589405

**Authors:** Este Leidmaa, Andreas Zimmer, Valentin Stein, Anne-Kathrin Gellner

## Abstract

**Background:** The blood-brain barrier (BBB) maintains brain homeostasis by protecting the brain from pathological stimuli and controlling the entry of physiological substances from the periphery. Consequently, alterations in BBB permeability may pose a threat to brain health. Long-term consumption of a high-fat high-sugar/Western diet (HFD) is known to induce BBB dysfunction. However, nothing is known about the immediate effects of acute HFD consumption on the BBB.

**Methods:** After consumption of either HFD or standard chow, mice were injected into the tail vein with fluorescent tracers of different sizes, including the drug doxorubicin. Individual brain regions were homogenized and analyzed for tracer extravasation using spectrophotometry. Localized tracer leakage over time in the somatosensory cortex was studied using high-resolution in vivo 2-photon microscopy.

**Results:** We demonstrate region-specific BBB leakage already after 1 hour of HFD for low- and high-molecular-weight tracers. Acute HFD also significantly increased BBB permeability to the anticancer drug doxorubicin.

**Conclusion:** These previously unknown effects of acute HFD may have direct and drastic implications for the clinical use of drugs depending on the dietary habits of the patient.

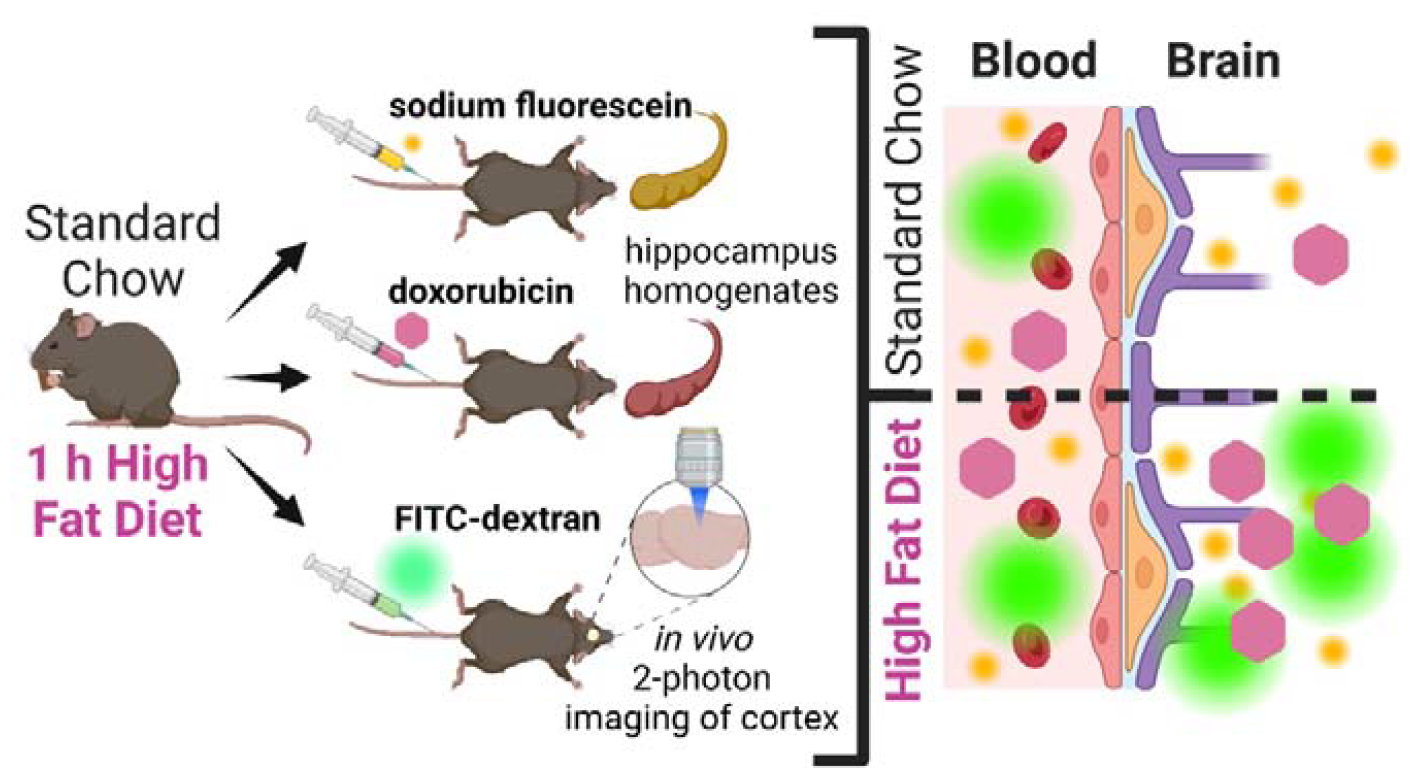

## Introduction

The blood-brain barrier (BBB) is a boundary between the bloodstream and the brain that tightly filters the passage of substances and nutrients from the blood to the brain. The BBB is clinically relevant in protecting the brain from toxins and pathogens, while maintaining a constant balance of water, nutrients, and hormone levels in the brain. The effective BBB also has a downside; many potential new drug candidates for mental and neurological diseases do not readily cross the BBB.

Alterations in BBB permeability can be induced by several physiological or pathological systemic and central states(1). In humans and mice alike, pathological triggers can be either short and strong, for example stroke, bacterial infection, trauma, or milder and longer-lasting like autoimmune disease, neurodegeneration, chronic stress, and obesity(2–8). Chronic consumption of a high-fat high-sugar/Western diet (HFD) can lead to obesity and related sequelae. Importantly, HFD has also been shown to induce BBB dysfunction(2,9). We were the first to show the effects of a very short HFD treatment of 1 hour and demonstrated that this acute HFD interfered with central mechanisms of feeding control(10). The impact of acute HFD on BBB permeability is completely unknown. Based on our previous observations, we now wondered whether such a short-term treatment would also affect BBB permeability. Surprisingly, using different methods, including *in vivo* microscopy, we demonstrate that the permeability of low- and high-molecular weight tracers, or the chemotherapeutic drug doxorubicin, is dramatically increased by this treatment.

## Materials and methods

### Mice and metabolic treatments

Adult male mice (C57BL/6N) were used for all experiments. Animals were single-housed throughout the *in vivo* imaging experiments starting after surgery, or for a minimum of 2 weeks prior to all other experiments. Mice were fed *ad libitum* and were housed under a 12:12-hour light/dark cycle (light phase from 7 am to 7 pm) at a constant temperature (22°C). Experiments were carried out in the beginning light phase. All experiments followed the guidelines of the German Animal Protection Law and have been approved by the government of North Rhine Westphalia (Local Committee for Animal Health, LANUV NRW).

Mice were weighed at start and the end of each treatment period. Mice were offered a free choice high-fat/high-sugar diet (HFD, EF 12451, ssniff Spezialdiäten, Germany) and standard chow (V-1534300, ssniff Spezialdiäten) or only standard chow for the control group for either 1 hour or 24 hours. HFD contained 4.615 kcal/g and had 45% of metabolizable energy from fat, 35% from carbohydrates (21% from sugar), and 20% from protein. Standard chow had 3.89 kcal/g and contained 9% of energy from fat, 58% from carbohydrate (4% of sugar), and 33% from protein. The amount of HFD consumed was measured. Of note, all mice chose to eat HFD and not standard chow during our treatment periods. To avoid neophobia, mice were offered one pellet of HFD 24 hours before the first treatment with HFD(10). Water was available *ad libitum* throughout the treatment periods.

### Sodium fluorescein assay

The protocol was modified from Kaya & Ahishali(11): Mice were anesthetized (medetomidine 0.5 mg/kg, midazolam 5 mg/kg bodyweight i.p.) after 1 hour of HFD or control treatment. Sodium fluorescein (F6377, Sigma) was dissolved in saline (30 mg/ml) and slowly injected into the tail vein according to body weight (120 mg/kg). After a circulation time of 30 min, mice were given a euthanizing dose of ketamine/xylazine (240 mg/kg, 32 mg/kg bodyweight i.p.). The mice were transcardially perfused with 50 ml cold phosphate-buffered saline (PBS, pH 7.4) at 10 ml/min. The brain was removed, and cortex, hippocampi, cerebellum, and hypothalamus were dissected, weighed, and snap frozen in liquid nitrogen. All further steps were carried out ensuring light protection of the samples. Tissue was thawn on ice, homogenized in 150 μl of PBS after which 150 μl of 60% trichloroacetic acid (T0699, Sigma) was added and the homogenates were vortexed for 2 min before incubation for 30 min at 4 °C. Samples were centrifuged at 18,000 g at 4°C for 10 min and 150 μl of supernatant per sample were measured for absorbance at 440 nm with a plate-reader (Infinite 200 Pro, Tecan) in a 96-well microplate. Tissue content of sodium fluorescein was quantified from a linear standard curve derived from the dye and expressed in nanogram per milligram of brain tissue.

### Cranial window surgery and 2-photon in vivo imaging

Mice were deeply anesthetized (medetomidine 0.5 mg/kg, midazolam 5 mg/kg, fentanyl 0.05 mg/kg body weight i.p.) and received carprofen 5 mg/kg s.c. for perioperative analgesia. A craniotomy 3-4 mm in diameter was carefully drilled over the left somatosensory cortex. A round glass coverslip 5 mm in diameter was placed and sealed with cyanoacrylate and dental acrylic over the craniotomy. A custom-made plastic bar was attached to the parietal bone contralateral to the trepanation for fixation during microscopy. Anesthesia was antagonized (atipamezole 2.5 mg/kg, flumazenil 0.5 mg/kg, naloxone 1.2 mg/kg body weight i.p.) and mice allowed to return to full alertness and normal mobility under a warming lamp. Carprofen (5 mg/kg body weight) was applied for analgesia 1x/day subcutaneously for 3 days starting 12 hours post-surgery.

After a minimum recovery period of 25 days mice were slightly anesthetized (medetomidine 0.5 mg/kg, midazolam 5 mg/kg body weight i.p., toe pinch slightly positive) to ensure immobilization during the imaging session. 70 kDa FITC-dextran (Sigma 4695/100mg/F) was dissolved in saline (50 mg/ml) and 100 μl were slowly injected into the tail vein. A custom-built 2-photon microscope equipped with a Chameleon Vision S laser (Coherent) and a water immersion objective lens (40x, NA 0.8, Olympus) was used for *in vivo* imaging. Images were acquired using ScanImage software (MBF Bioscience). Excitation wavelength was tuned to 910 nm for imaging of FITC-Dextran. Time-lapse stacks (xyz-dimension 334 x 334 x 40 μm, xy-resolution 0.33 μm / px, step size 2 μm) of a region of interest (ROI) containing a vascular network were recorded every 3 minutes for 30 minutes at a frame rate of 0.43 Hz. After imaging, anesthesia was antagonized with atipamezol 2.5 mg/kg and flumazenil 0.5 mg/kg body weight i.p and mice were allowed to fully recover in their home cage under a warming lamp. Imaging sessions of the same animal were conducted with a minimum interval of 5 and a maximum interval of 12 days. Treatments before the imaging sessions were done in a pseudorandomized order and the area with the region of interest was re-identified each time using a 4x objective. By this, every animal served as its own control except for one which accidentally lost its cranial window before the standard chow condition. The results of one mouse after 1 hour of HFD had to be excluded as a technical outlier.

### Image analysis

Preprocessing and analysis of the *in vivo* 2-photon images were done with Fiji/ImageJ2 (2.9.0). The z-stacks from all the time-points were transformed into maximal projections which were then added together into a time-stack (10 frames in 3-minute steps). Image registration was done with “Linear Stack Alignment with SIFT” and expected transformation “Translation”. Four rectangular regions of interest (ROIs) of 16.5 x 16.5 μm were placed to the intravascular space with at least a 2 μm distance from vessels and 4 ROIs of 33 x 33 μm were placed on the vessels as a control measure (Figure 1F). Fluorescence intensity (mean gray value) was measured for all ROIs. Data from the 4 extravascular and 4 intravascular ROIs were averaged respectively per imaging session into one set of time-lapse values. All the subsequent time points were normalized to the first time point (T_n_-T_0_), which served as the baseline. Areas under the curve (AUC) were additionally calculated for each imaging session from every animal using GraphPad Prism (Version 9.4.1).

**Figure 1.**
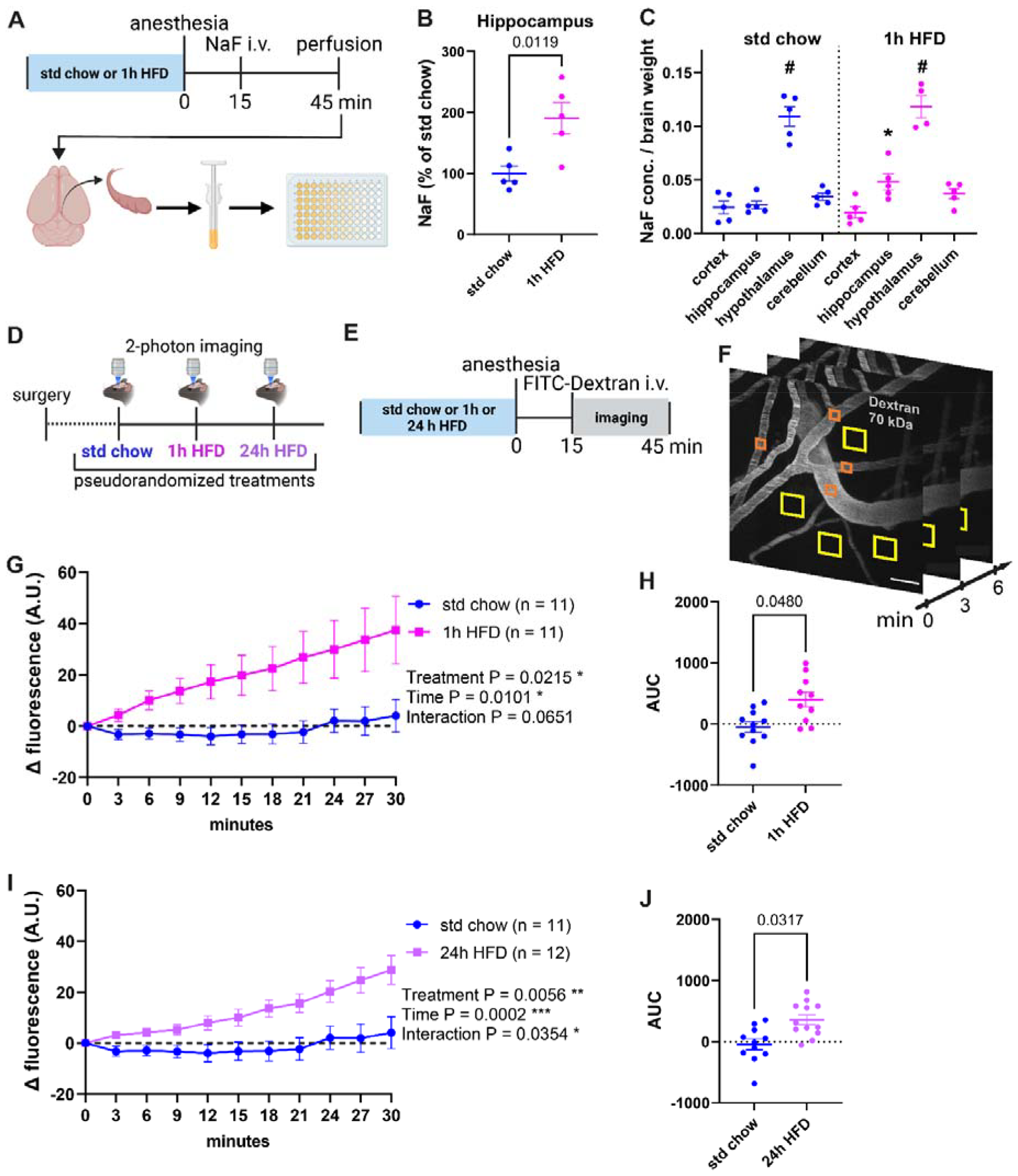
Leakage of the small-molecular-weight tracer sodium fluorescein (NaF, 376 Da) and a high-molecular-weight tracer FITC-dextran (70kDa) from blood into the brain tissue was increased already after 1 hour of high-fat high-sugar diet (HFD) exposure. **(A)** Scheme showing the design of the experiment with NaF and workflow of the tissue processing with dissection of the regions of interest, homogenization, and fluorometric measurements. **(B)** Relative NaF concentrations in hippocampal homogenates were elevated (t(8) = 2.58, p = 0.0327) after 1h of HFD compared to control mice on standard chow. **(C)** The hypothalamus showed the highest absolute NaF leakage into the brain tissue compared to the other regions both in control condition (F(1.561, 8.325) = 46.08, p < 0.0001) and after 1h HFD (F(1.428, 5.236) = 47.53, p = 0.0006). P < 0.05, * hippocampus compared to cortex,#hypothalamus compared to all other brain areas. Error bars represent mean ± SEM. **(D)** Scheme showing the design of the experiment with repeated *in vivo* imaging of dextran-filled cortical vasculature and perivascular space in the somatosensory cortex (S1) after 3 pseudorandomized treatments of either standard chow, 1 hour, or 24 hour HFD. **(E)** Workflow of treatment and dextran injection before an imaging session. **(F)** Example of dextran-filled vasculature and analysis of intra- (orange squares) and extravascular (yellow squares) fluorescence in maximum projections of time-lapse z-stacks, scale bar 50 μm. **(G)** Acute 1h HFD significantly increases extravascular FITC-dextran levels in the somatosensory cortex over the time course of imaging (treatment F(1, 11) = 7.173, p = 0.0215, time F(1.525, 16.78) = 6.897, p = 0.0101, time x treatment interaction F(1.241, 10.92) = 3.989, p = 0.0651) and the leakage was also significantly higher (t(8) = 2.33, p = 0.0480) when expressed as area under the curve (AUC) **(H). (I)** Exposure to HFD for 24h increased the leakage of FITC-dextran into the somatosensory cortex significantly for analysis over time (treatment F(1, 11) = 11.81, p = 0.0056, time F(2.407, 26.47) = 10.97, p = 0.0002, time x treatment interaction F(1.53, 15.14) = 4.593, p = 0.0354) and when expressed as AUC (t(10) = 2.49, p = 0.0317) **(J)**. Error bars represent mean ± SEM.

### Doxorubicin assay

The protocol was modified from Bachur and colleagues(12): Mice were anesthetized (medetomidine 0.5 mg/kg, midazolam 5 mg/kg body weight i.p.) after 1 hour of HFD or control treatment. Doxorubicin-hydrochloride (44583, Sigma) was dissolved in saline (1.25 mg/ml) and slowly injected into the tail vein according to body weight (5 mg/kg). After a circulation time of 60 min, at which peak concentrations of doxorubicin can be expected(13) mice were given a euthanizing dose of ketamine/xylazine (240 mg/kg, 32 mg/kg body weight i.p.). The mice were transcardially perfused with 50 ml cold phosphate-buffered saline (PBS, pH 7.4) at 10 ml/min. The brain was removed, and cortex, hippocampi, cerebellum, and hypothalamus were dissected, weighed, and directly homogenized in 10 volumes of 0.3N HCl in 50 % ethanol. All further steps were carried out ensuring light protection of the samples. Homogenates were incubated at 4°C for 24 hours before centrifugation at 16,000 g for 25 min at 4°C. Supernatants were collected and 100 μl of each sample measured with a spectrophotometer (Nanodrop 2000c, Thermo Scientific). Doxorubicin concentrations were determined from absorbance at 480 nm and a linear standard curve derived from the compound and expressed in nanograms per gram of brain tissue.

### Statistics

Data was analyzed and plotted with GraphPad Prism, Versions 9.4.1. and 9.5.0. Drawings were created with BioRender.com. The numbers of animals/samples are indicated in the legends. Statistical outliers were identified using the ROUT method (Q =1 %). The Shapiro-Wilk test was used to estimate whether the data was normally distributed. Statistical significance was assessed using Student’s t-test for parametric data or Mann– Whitney test for nonparametric data. One-way ANOVA was used for matched values/Mixed model analysis with Geisser-Greenhouse correction and Tukey’s multiple comparison test was used to analyze more than two groups. Repeated measures 2-way ANOVA/Mixed model analysis with Geisser-Greenhouse correction with Sidak’s multiple comparison tests were used to evaluate time-lapse values.

## Results

We injected the low molecular weight tracer sodium fluorescein (376 Da) intravenously in anesthetized mice after they had eaten HFD for 1 hour and subsequently analyzed the levels of sodium fluorescein in brain homogenates (Fig. 1A). As shown in Fig. 1B, HFD exposure resulted in a significant increase in tracer fluorescence specifically in the hippocampus, thus indicating that the BBB permeability was increased after acute HFD. No acute effect of HFD was detectable in samples from cortex, cerebellum, and hypothalamus by this method (SupplFig. 1B-D). Of note, the hypothalamus with its specialized fenestrated BBB(14) showed markedly higher fluorescence levels compared to the other brain regions in both treatment groups (Fig. 1C).

We next investigated BBB leakage of the high molecular weight tracer FITC-dextran (70 kDa) using 2-photon *in vivo* imaging through a cranial window over the somatosensory cortex after intravenous injection in anesthetized mice (Fig. 1D-F). The somatosensory cortex is known to be activated by the perception and processing of stimuli such as palatable food in humans(15,16). This *in* vivo imaging method provides a high spatial resolution, allows for repeated measurements over several time points in each individual, and enables within-subject comparison. First, we measured the effect of 1 hour of HFD consumption versus standard chow within 30 minutes after tracer injection. Whereas mice fed with a standard chow showed no increase of extravascular FITC-dextran during the 30-minute observation period, we detected a significant leakage of FITC-dextran into the extravascular space in mice exposed to HFD. The control measure of intravascular fluorescence decreased over time as expected (SupplFig. 1E). We repeated the analysis after an exposure to a 24-hour HFD treatment, which is known to increase BBB leakage in the hippocampus(17). Strikingly, the 24-hour treatment increased the BBB leakage to a similar extent as the 1-hour treatment when compared over the time course (Fig. 1G, I) and when the cumulative areas under the curve (AUC) were analyzed (Fig. 1H, J). Direct comparison also showed no significant difference between the two treatment groups (SupplFig. 1F). Together, these findings indicate that the 1-hour HFD already produced a comparable effect to that of a prolonged treatment.

We next sought to determine if the unexpectedly rapid modulation of the BBB function could be clinically relevant. For this purpose, we tested the BBB penetrance of doxorubicin-hydrochloride, a widely used chemotherapeutic agent which is effective against various cancers(18). Doxorubicin-hydrochloride has a relatively poor BBB penetrance, which is an advantage in the treatment of non-brain tumors. For brain tumors, however, an improved BBB permeability would be advantageous, and this has been the aim of different studies(13,19). The intrinsic fluorescence of doxorubicin can be used as a proxy for its entry to the tissue. When we injected doxorubicin-hydrochloride (580 Da) intravenously after 1 hour of HFD exposure (Fig. 2A), fluorescence was significantly increased in hippocampal homogenates 1 hour after treatment compared to the standard chow condition. (Fig. 2A-B). As we had previously observed for sodium fluorescein (376 Da), no differences were detected in the cortex, cerebellum, and hypothalamus (SupplFig. 2B-D). Again, the hypothalamus showed higher fluorescence levels compared to the other brain regions in both treatment groups, while the increase seen in the cerebellum was unique to doxorubicin (Fig. 2C).

**Figure 2.**
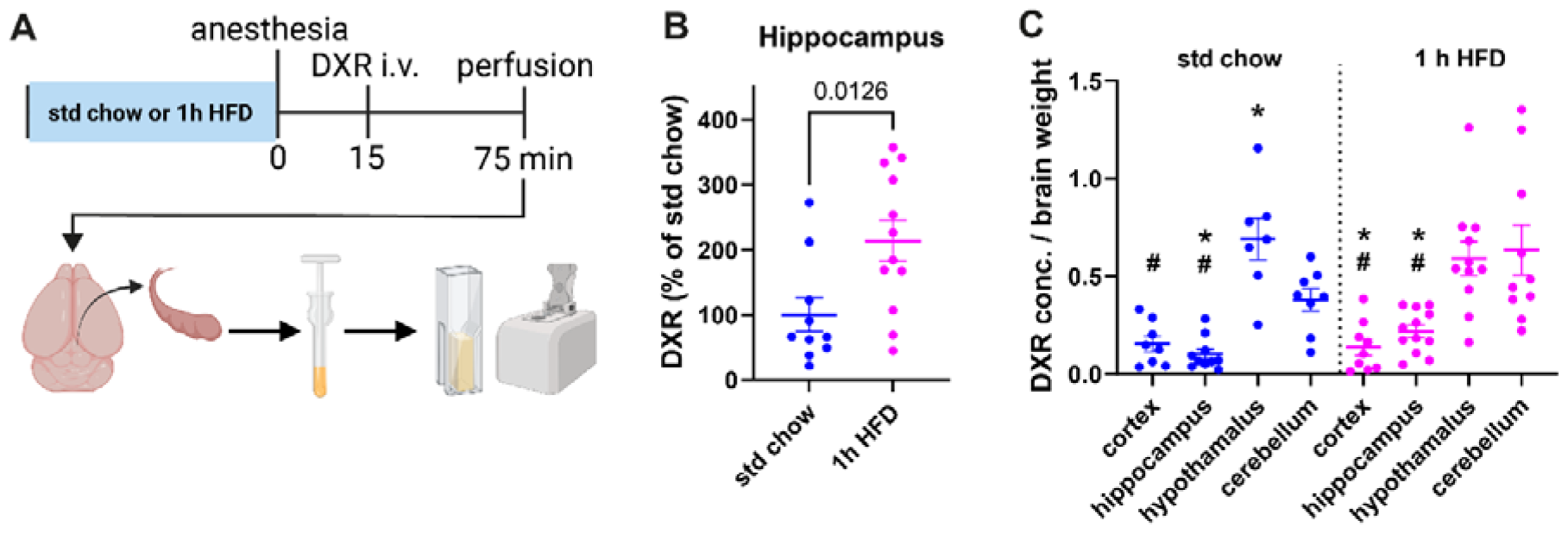
Leakage of the chemotherapeutic agent doxorubicin (DXR, 580 Da) from blood into the brain tissue was increased in the hippocampus after 1 hour of high-fat high-sugar diet (HFD) exposure. **(A)** Scheme showing the design of the experiment and workflow of the tissue processing with dissection of the regions of interest, homogenization, and spectrophotometric measurements. **(B)** DXR was elevated in the hippocampus (t(20) = 2.74, p = 0.0126) after 1 hour of HFD normalized to control mice on standard chow. **(C)** Hypothalamus and cerebellum showed the highest absolute NaF leakage into the brain tissue compared to the other regions both in control condition (F (3, 29) = 20.21, p <0.0001) and after 1h HFD (F (3, 38) = 9.911, p <0.0001). P < 0.05, * compared to cerebellum,#hypothalamus compared to all other brain areas. Error bars represent mean ± SEM.

## Discussion

In this paper, we demonstrate an increased BBB permeability after acute exposure to HFD, by analyzing the uptake of low-molecular-weight tracers in hippocampal homogenates, as well as the leakage of a high-molecular-weight tracer into the cortex using 2-photon *in vivo* microscopy. The magnitude of BBB permeability was similar after a 1-hour treatment and a 24-hour exposure, indicating that an acute treatment already produced a maximal effect. Our findings might be of drastic clinical relevance because a 1-hour treatment also increased the BBB permeability for the chemotherapeutic drug doxorubicin. This finding indicates that certain pharmacotherapies may be acutely affected by dietary factors in unexpected ways.

Many studies have addressed changes in BBB permeability under pathological conditions but also upon physiological stimuli, including high-caloric sugar and fat-containing diets(1,20). This is particularly important when considering the global spread of such an unhealthy diet. Indeed, several studies demonstrated that longer periods of HFD exposure, 24 hours and more, increased tracer penetration into brain tissue(2,9,17). Unfortunately, the molecular and cellular mechanisms underlying the effects of HFD on the BBB remain elusive.

The fact that we detected rapid leakage in hippocampal homogenates but not in other brain regions led to using a method with much higher spatial resolution. Two-photon *in vivo* microscopy allows repeated analysis of the cortical BBB in different conditions, as we have previously demonstrated(21). Indeed, with this high-resolution method, we detected rapid and robustly increased extravasation of a high-molecular-weight tracer as large as plasma albumin (70 kDa) into the somatosensory cortex after 1 and 24 hours of HFD. Since there was no detectable dose-dependency regarding treatment duration, we assume that the naturally intermittent diurnal feeding behavior sustains BBB leakage upon HFD exposure of more than 1 hour. Whether BBB permeability further increases with HFD treatment longer than 24 hours remains to be elucidated with the *in vivo* imaging approach.

It is important to note that volatile agents such as iso- and sevoflurane also disrupt the BBB(22). For this reason, we have used an injectable anesthetic cocktail consisting of medetomidine, midazolam, and fentanyl in our study. Administration of dexmedetomidine, an active metabolite of medetomidine, has been shown to prevent BBB disruption(23) and repeated injection of midazolam also has protective effects against BBB leakage(24). Thus, our results are even likely to underestimate the effects of acute HFD.

Putative mechanisms of rapid HFD-induced BBB leakage could be hypothesized based on experiments with short-term treatment periods: an increase in inflammatory markers in the murine hippocampus and hypothalamus were reported after 48 and 24 hours of HFD, respectively(17,25). However, many regulators of BBB permeability have been identified in addition to inflammatory cytokines, including nitrous oxide, calcium influx, or hemodynamic changes(1). Overall, the dynamic modulation of the BBB could also serve a physiological purpose, for example allowing faster transport of peptide hormones that regulate feeding and nutrient metabolism into the brain(20,26). With this in mind, acute HFD could also influence BBB permeability through novel and specific mechanisms. Thus, several possible mediators of the rapid HFD-induced BBB alteration have to be explored next.

Finally, the striking finding that acute HFD-induced BBB leakiness for two exogenous molecules of different sizes led to the clinically relevant question of whether acute dietary choices could also impact the brain delivery of drugs that do not penetrate the BBB well. We demonstrated a 2-fold increased concentration of the chemotherapeutic agent doxorubicin in the hippocampus after eating HFD for 1 hour. This should be considered in the context of drug treatments: if the target is a brain tumor, eating high amounts of energy-dense food could be used to enhance drug delivery. On the other hand, increased concentrations of a neurotoxic drug in healthy brain tissue could lead to severe side effects. Here, we provide the foundation for an entirely new line of research that considers how acute dietary stimuli modulate BBB permeability in pharmacotherapy.

## Declarations

### Availability of data and materials

All requests should be sent to the corresponding authors AKG or EL. The datasets generated during and/or analyzed during the current study are available from the corresponding authors on reasonable request.

### Authors’ Contributions

Conceptualization: EL, AZ, VS, and AKG; Methodology: EL and AKG; Investigation: EL and AKG; Writing—original draft: EL and AKG; Writing—review and editing: EL, AZ, VS, and AKG; Funding acquisition: EL and AKG; Supervision: EL and AKG. All authors contributed to the final version of the manuscript.

