## Supplementary figures for "Acute high-fat high-sugar diet rapidly increases blood-brain barrier permeability in mice"

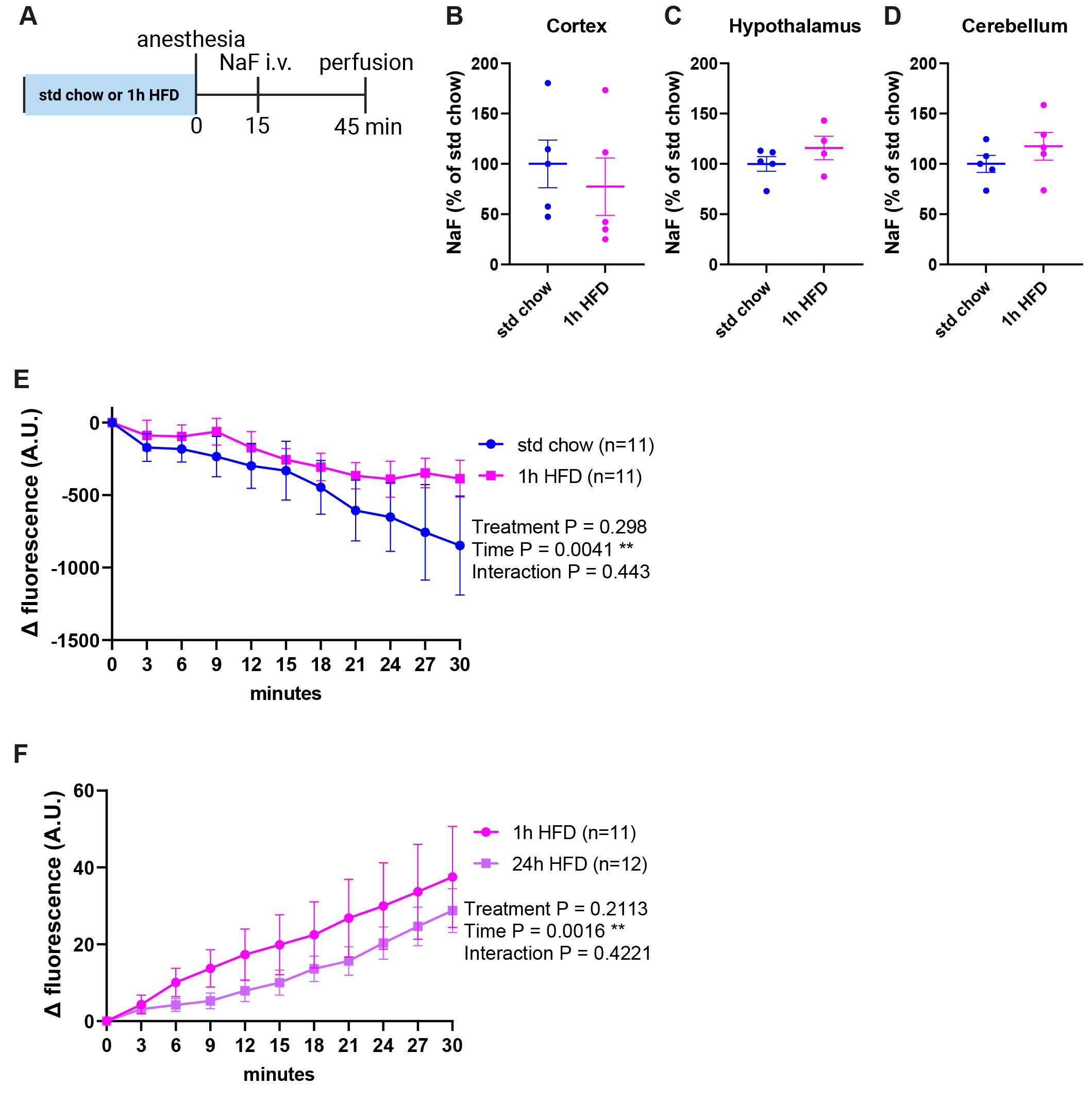

**Supplementary Figure 1. Unaltered leakage of small molecular weight tracer sodium fluorescein (NaF) (376 Da) from blood into the brain tissue in cortex, hypothalamus, and cerebellum after 1 hour of high-fat high-sugar diet (HFD) exposure.**

**(A)** Scheme showing the design of the experiment. Relative NaF concentrations were not altered (t(8) = 0.610, p = 0.559) after 1h of HFD in cortex **(B)**, hypothalamus (t(7) = 1.22, p = 0.263) **(C),** or cerebellum (t(8) = 1.09, p = 0.308) **(D)**. Changes of intravascular FITC-dextran levels in the somatosensory cortex are significant over the time course of imaging but do not differ by treatment; treatment F(1, 11) = 1.95, p = 0.298, time F(2.1, 23.1) = 6.866, p = 0.0041, time x treatment interaction F(1.922, 16.91) = 0.844, p = 0.443 **(E)**. The leakage of FITC-dextran into the somatosensory cortex is not statistically different when comparing 1- and 24-hour exposure to HFD; treatment F(1, 11) = 1.762, p = 0.211, time F(1.167, 12.83) = 14.6, p = 0.0016, time x treatment interaction F(1.508, 14.93) = 0.835, p = 0.422 **(F)**. Error bars represent mean ± SEM.

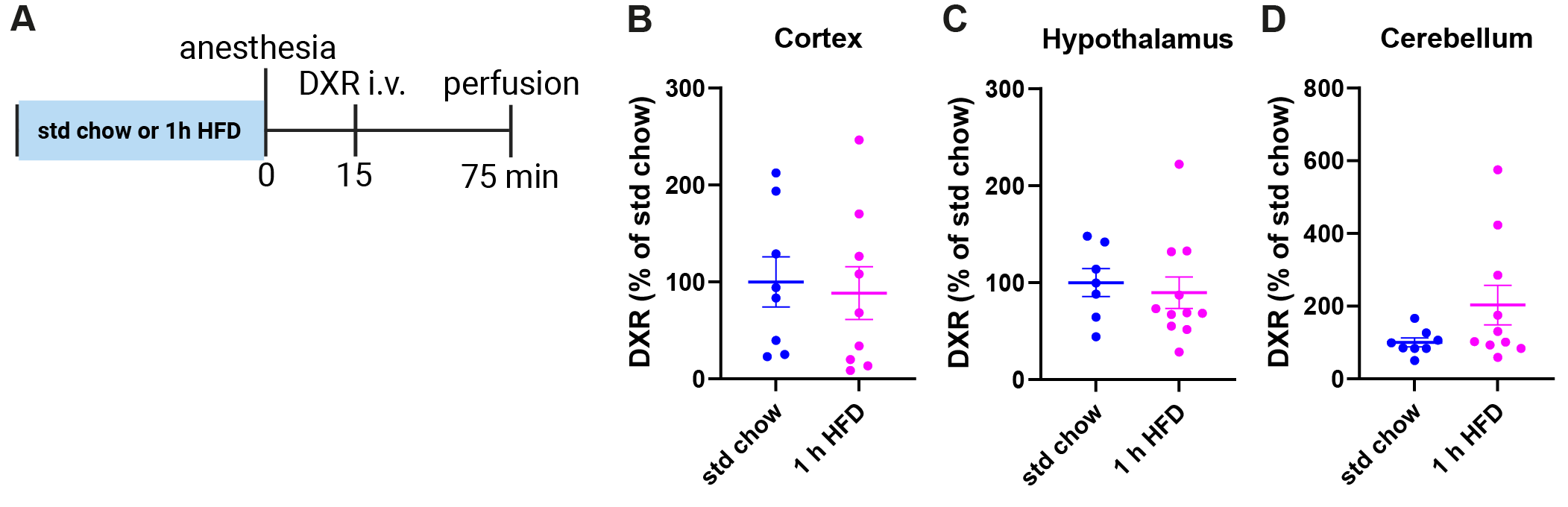

**Supplementary Figure 2. Unaltered leakage of doxorubicin (580 Da) from blood into the brain tissue of cortex, hypothalamus, and cerebellum after 1 hour of high-fat high-sugar diet (HFD) exposure.**

**(A)** Scheme showing the design of the experiment. Relative DXR concentrations were not altered after 1 hour of HFD in cortex (t(15) = 0.309, p = 0.762) **(B)**, hypothalamus (U = 29, p = 0.425) **(C),** or cerebellum (U = 23, p = 0.146) **(D)**. Error bars represent mean ± SEM.
